# Multiphoton imaging of glucose, galactose, and fructose-induced formation of fluorescent advanced glycation end products in tissues

**DOI:** 10.1101/2023.01.04.522782

**Authors:** Chih-Ju Lin, Sohidul Mondal, Sheng-Lin Lee, Jeon-Woong Kang, Peter T. C. So, Chen-Yuan Dong

## Abstract

Blood glucose and HbA1c, intermediate glycation products of hemoglobin, remain the two clinical biomarkers for monitoring disease progression in diabetics. However, the formation of advanced glycation end products (AGEs) has been implicated in diabetic pathogenesis and the use of AGEs in tissues as long-term glycemic markers may be of value in the clinical setting. Therefore, it is necessary to understand how different tissue constituents respond to dietary monosaccharides. In this study, we studied the *in vitro* rate of fluorescent AGEs (fAGEs) formation with multiphoton microscopy in different porcine tissues (aorta, cornea, kidney, dermis, and tendon). These tissues were treated with D-glucose, D-galactose, and D-fructose, three primary monosaccharides found in human diets. We found that the use of D-fructose resulted in the highest glycation rate, followed by D-galactose and then D-glucose. Moreover, compared to non-collagen tissue constituents such as elastic fibers and cells, the rate of tissue glycation was consistently higher in collagen, suggesting that collagen is a more sensitive target for fAGE formation. However, we also found that collagen in different tissues exhibits different rates of fAGE formation, with slower rates observed in tightly packed tissues such as cornea and tendon. Our study suggests that for fAGE to be developed into a long-term glycemic biomarker, loosely organized collagen tissues located in the proximity of vasculature may be the best targets.

## Introduction

Diabetes mellitus is a worldwide health epidemic. With an estimated prevalence of 415 million in 2015, diabetes affects the health of 8.8% of the global population between the age of 20-79. With a projected prevalence of 642 million in 2040, diabetes will be an important health issue for many years to come. Since diabetes is a long-term medical condition, the cost of caring ($673 billion in 2015) for affected patients is staggering [1]. In diabetic patients, the presence of elevated sugar levels leads to a non-enzymatic Maillard reaction with amino groups, first leading to the formation of Schiff bases which are later converted to Amadori products [2]. While these are reversible intermediate glycation products, they eventually convert to long-lived and irreversible advanced glycation end products (AGE). While tissue glycation occurs with normal aging, AGE accumulation is accelerated in diabetic patients. AGEs affect normal physiological function by forming additional cross-links in connective tissues, and alteration of cellular function by binding to receptors for AGE (RAGE) which can result in the production of reactive oxygen species [3–5]. AGE accumulation in tissues has been implicated in diabetic complications such as retinopathy, corneal alterations, nephropathy, neuropathy, and cardiovascular diseases [6–11]. The roles of AGEs in diabetic pathogenesis also led to concerns about the diet input of AGEs as the Maillard reaction is accelerated at higher temperatures and leads to food browning [12, 13]. In fact, feeding laboratory animals with AGEs can lead to atherosclerosis and kidney diseases [14, 15].

The current approach to managing diabetic patients is largely based on the Diabetes Control and Complications Trial (DCCT) study from 1983-1993. This seminal study demonstrated that the effective control of blood glucose levels can reduce the effects of diabetic complications in retinopathy, nephropathy, and neuropathy [16]. As a result, the frequent measurement of blood glucose levels and periodic determination of glycated hemoglobin HbA1c became standard clinical practice in managing diabetic patients. However, both blood glucose and HbA1c measurements are transient and do not reflect the cumulative change to connective tissues caused by prolonged exposure to elevated blood sugar levels.

In the past, AGE inhibitors such as aminoguanidine are effective in reducing the pathogenesis of diabetic complications [9, 17]. Other molecules such as 3-phenyacyl-4,5-dimethylthiazolium chloride can improve compliance in stiffened arteries [18, 19]. Due to the importance of AGE inhibition, various strategies, including the use of various plant extracts, have been investigated [20–22]. The DCCT Skin Collagen Ancillary Study Group showed that corrected for HbA1c, AGE level from skin collagen is most consistent with diabetic complications and that AGE detection can be developed into a diagnostic tool that may surpass the diagnostic value of HbA1c measurement [23]. In fact, among the variety of AGE species, some, such as pentosidine are known to be fluorescent [21, 22, 24–26].

Several studies have demonstrated the correlation between tissue fluorescence and diabetes. In the case of skin, it was shown that adjusted for age, collagen from skin biopsy contains a significantly higher level of fluorescence in diabetics. In addition, it was found that the level of fluorescence increases with the severity of retinopathy, nephropathy, and stiffness of the artery and joints [27]. These studies eventually led to the development of non-invasive measurement of skin and ocular lens fluorescence as a tool for diagnosing diabetic complications. Increased skin fluorescence correlated with age, diabetic duration, HbA1c from the previous year, and creatinine level [28, 29]. In the case of a lens, it was found that corrected for age, fluorescence increases from normal, prediabetics, type II diabetics, and type I diabetics [30]. In the case of skin, substantial clinical data has been accumulated correlating skin fluorescence in diabetics and non-diabetics. These studies include kidney disease [31], microvascular/macrovascular complications [32], cardiovascular pathology [33], and peripheral vascular systems [34]. In all cases, while a difference in skin fluorescence was found between diabetic and non-diabetic patients, a substantial overlap in fluorescence intensity was found between the two groups. This observation may be interpreted by the presence of other intrinsic fluorescent species such as NADH and flavins [35–37]. Differences in skin tone which can affect the penetration of excitation wavelength and detection of fAGEs, and differences in individuals’ sun exposure which may lead to fAGE photobleaching. Finally, an assay for diabetic complications based on fAGE can be complicated by differences in sugar consumption. It is known that the rate of glycation is different for different monosaccharides, with aldoses, in general, being more reactive than ketoses. For commonly simple sugars found in human diets, sugar reactivity with the amino group in forming Schiff’s base is found to be in the order of glucose, galactose, and fructose [38]. With a change in the modern diet and the associated increase in the use of fructose in processed foods, a clarification of the fluorescence properties of fAGE formed from different monosaccharides may be beneficial in the development of a technology that uses fAGE as a diabetic biomarker. Moreover, as different tissues may react differently to glycation, there is a need to study the rate of fAGE formation in organs that may be affected by diabetes.

## Materials and Methods

### Porcine tissues preparation

In this study, we chose to study the rate of fAGE formation in the porcine aorta, cornea, kidney, skin dermis, and tendon. Aorta and kidney were chosen as they represent organs that can be seriously affected by diabetes. Cornea, skin, and tendon were chosen as these tissues may be accessed for predictive evaluation of diabetic pathogenesis in internal organs. To prepare the aorta specimens, we acquired a porcine heart to remove descending aorta located approximately 10-15 cm from the heart. We processed the aorta such that each blood vessel was approximately 2 cm in outer diameter and 3 cm in length. The artery was then processed into sections 2 mm in thickness before being immersed in the incubation solutions. To obtain the cornea specimens, porcine eyes were acquired and the epithelium layer was scraped. Next, an 8 mm punch (Harris Uni-Core, Hatfield, PA) was used to excise the porcine cornea specimen. Kidney tissues were acquired and the renal cortex was processed into sections each approximately 2 mm in thickness and 5×5 mm^2^ in area. Skin dermis tissues were acquired from porcine dorsal skin. After removing the fat layers, the exposed dermis was cut into sections with a volume of about 2×6×5 mm^3^. Tendon tissue was acquired from the porcine patellar tendon. Since the patellar tendon has muscle tissues on both ends, we only used the middle sections which were further cut into strips 10 mm in length, 2 mm in thickness, and 6 mm in width. Prior to immersion in the glycation solutions, the tissue sections were sterilized with 4% proviodine-iodine solution for 30 s and rinsed 3 times in a sterile 1× phosphate-buffered saline (PBS) solution. For glycation incubation, two pieces of each tissue type were immersed for each incubation condition.

### Tissue incubation

For this study, we used three primary monosaccharides found in the human diet: D-glucose (Sigma-Aldrich, St. Louis, MO), D-fructose (Sigma-Aldrich, St. Louis, MO), and D-galactose (Acros Organics, Fair Lawn, NJ). Since disaccharides and polysaccharides such as sucrose, lactose, and starch require enzymatic digestion into monosaccharides before intestinal absorption [33], we will focus on monosaccharides in this study. Each monosaccharide was prepared into a concentration of 0.5 M in 0.05 M PBS (phosphate-buffered saline) solutions. In addition, 1% penicillin-streptomycin (10,000 U/mL, Gibco, Waltham, MA) was added to inhibit bacterial growth. The tissue sections were independently immersed in 15 mL centrifuge tubes, each centrifuge tube was then filled with 5 mL incubation solution for complete immersion of the tissue sections. The specimens were then incubated in a 37 °C incubator, with the incubation solutions, replaced every 4 days. Image acquisition was performed every 8 days. Glycated tissue sections were placed onto glass slides and covered with No. 1.5 cover glasses.

### Multiphoton microscopy

Multiphoton imaging was performed on a home-built system based on an inverted microscope (TE2000U, Nikon, Japan). The excitation source was a titanium-sapphire (ti-sa) laser (Tsunami®, Spectra Physics, Santa Clara, CA) pumped by a diode-pumped, solid-state laser operating at 532 nm (Millennia^®^ Pro, Spectra Physics, Santa Clara, CA). 780 nm output of the ti-sa laser was used as the excitation source. Upon reflection from a galvanometer-driven, x-y scanning system (6215M, Cambridge Technology, Watertown, MA), the laser was beam-expanded and reflected into the focusing objective (20X, S Fluor, 20×/NA 0.75, Nikon, Japan) by a primary dichroic mirror (720dcspxr, Chroma Technology, Bellows Falls, VT). Second harmonic generation (SHG) and specimen autofluorescence were collected for the focusing objective in the epi-illuminated geometry. After short-pass filtering by the primary dichroic, additional dichroic mirrors (420dclr, 495dcxr, and 550 dcxr, Semrock, Rochester, NY), and bandpass filters (FF01-390/18, Semrock; HQ460/50, Chroma Technology; FF03-525/50, FF01-630/92, Semrock). Luminescent signals from the specimen were separated into SHG (390 ± 9 nm), blue (460 ± 25 nm), green (525 ± 25 nm), and red (630 ± 46 nm) fluorescence for detection. Single photon counting photomultipliers (PMT, R7400P, Hamamatsu, Hamamatsu City, Japan) were used to detect signal photons.

The average laser power on the sample was 30 mW, the laser beam was focused on the tissue and the scanned area was 256 × 256 pixels in size, corresponding to an area of about 230 × 230 μm^2^. We randomly selected five ROIs for imaging of the cornea, tendon, aorta, skin, and kidney samples. Aortic images were obtained near the inner rings of the aorta samples which contain endothelium, tunica intima, and media layers. We focused on imaging the tunica intima and media layers which are largely composed of collagen and elastic fibers. For skin imaging, we focused on the dermal layers, and the epidermis was avoided. For each sample, five locations are chosen for analysis of the collagen fluorescence using SHG imaging to locate collagen fiber. Following the selection of the regions, we analyze the fluorescence intensity by reading the same ROIs on fluorescence imaging.

## Results and Discussion

An earlier study measuring *en face* fluorescence of the skin demonstrated a marked increase in the 430-590 nm range [24]. Our study found that ribose-induced fAGEs in collagen have fluorescence emissions that peak at around 500 nm [31, 32]. These results suggest that AGEs with different fluorescence spectral properties may be produced by reacting with different sugar molecules in tissues.

After performing multiphoton imaging of the incubated tissue specimens, we merged the red, green, and blue range fluorescent signals to a single autofluorescence channel, separated from collagen SHG. Time-lapsed two-photon autofluorescence (TPAF) and SHG images of glycated and control tissue sections for porcine aorta (Fig. 1), the cornea (Fig. 2), kidney (Fig. 3), dermis (Fig. 4), and tendon (Fig. 5) were acquired. In each data set, we show images of the tissue sections incubated in control, D-glucose, D-galactose, and D-fructose solutions after 0, 16, 24, 32, and 40 days. Since specimens on Day 8 have not been significantly glycated, images at that time point are not shown. As Fig. 1 shows, at Day 0, both collagen and elastic fibers are present. Note that in control samples, elastic fibers can be visualized by the presence of their autofluorescence and the absence of SHG signals while collagen fibers produce strong SHG signals. With the increase in incubation time, we notice an increase in autofluorescence intensity of the aorta specimens incubated in the monosaccharide solutions, with the most substantial increase observed in the D-fructose-immersed specimens. Qualitatively, D-glucose- and D-galactose incubated aorta sections also showed an increase of autofluorescence, but at a significantly lower rate than that due to D-fructose. Similarly, Fig. 2 shows autofluorescence images of the incubated cornea specimens. As cornea stroma is composed primarily of collagen, its presence can be detected through SHG signals. Similar to the aorta results, fAGE formation is most significant in D-fructose-treated specimens, followed by D-galactose and D-glucose-treated specimens. In the case of the renal capsule (Fig. 3), similar trends are seen in the glomeruli and tubules where there is a marked increase in the tubular endothelial and glomerular cells. Since adverse effects on renal functions due to diabetes contribute to glomerular hyperfiltration, therefore, increased fAGE levels in cells of the renal cortex may be indicative of diabetic pathogenesis. We also studied the temporal dependence of fAGE formation in glycated skin dermis (Fig. 4). As the skin dermis is composed of collagen and autofluorescent elastic fibers. SHG and autofluorescence may be used to identify the presence of these fibers. In both tissue types, dermal collagen and elastic fibers demonstrate fAGE production following incubation in a monosaccharide solution, again, with D-fructose showing the largest increase in the rate of fAGE formation. The last tissue system we studied was the bovine tendon. Fig. 5 shows that regardless of the choice of incubation solution, fAGE formation was not apparent within 16 days of incubation. As glycation progresses, tendon autofluorescence started to increase with time. From the images, we further analyzed the rate of fAGE formation in different tissue constituents such as collagen, elastic fibers, and other organelles. Since the SHG signal can be used to identify collagen fibers, autofluorescence analysis can be performed on collagen fibers and other organelles (elastic fibers and cells). In the case of cornea and tendon, the tissues are composed primarily of collagen and its autofluorescence was used to analyze fAGE formation in collagen. However, in the skin dermis and aorta, fAGE autofluorescence is collected from collagen, elastic fibers, and cells.

**Figure 1.**
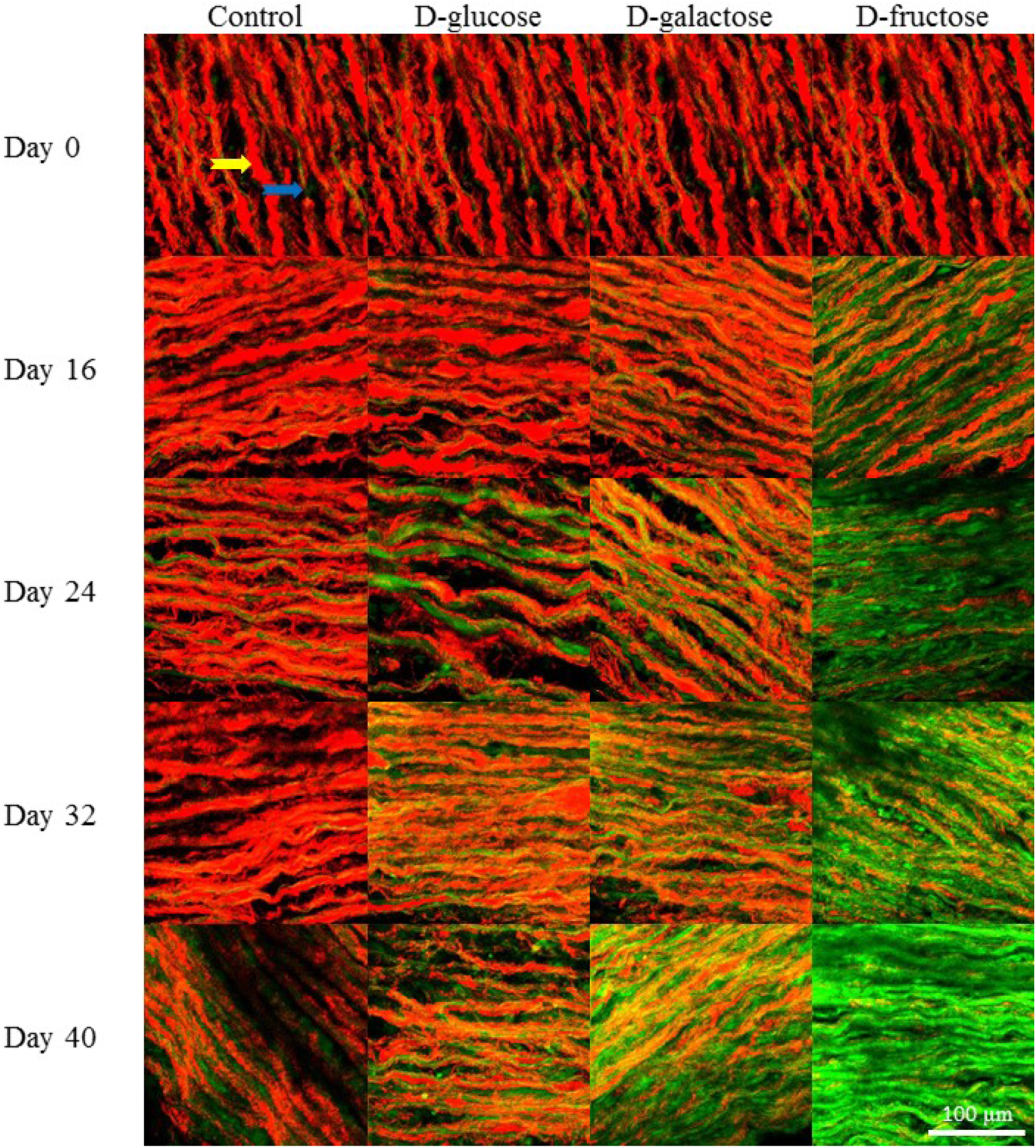
Autofluorescence and SHG imaging of porcine aorta sections that have been incubated in control, D-glucose, D-galactose and D-fructose solutions. Collagen (yellow arrow) is the main component of intima and media of aorta. Also present are autofluorescent elastic fiber (blue arrow). Red: SHG. Green: autofluorescence. Scale bar: 100 μm.

**Figure 2.**
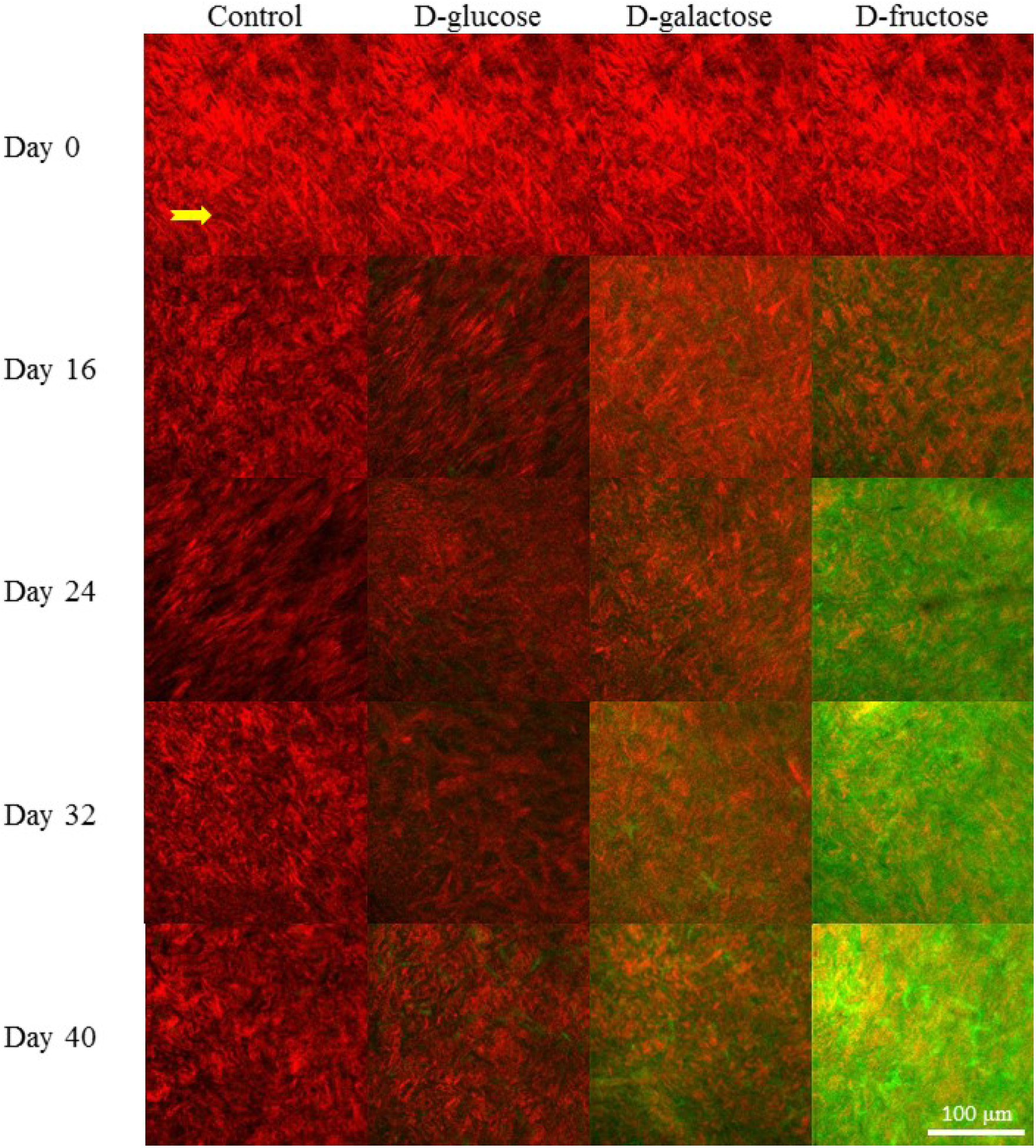
Autofluorescnece and SHG imaging of porcine cornea sections that have been incubated in control, D-glucose, D-galactose and D-fructose solutions. Collagen (yellow arrow) is the main component of cornea. Red: SHG. Green: autofluorescence. Scale bar: 100 μm.

**Figure 3.**
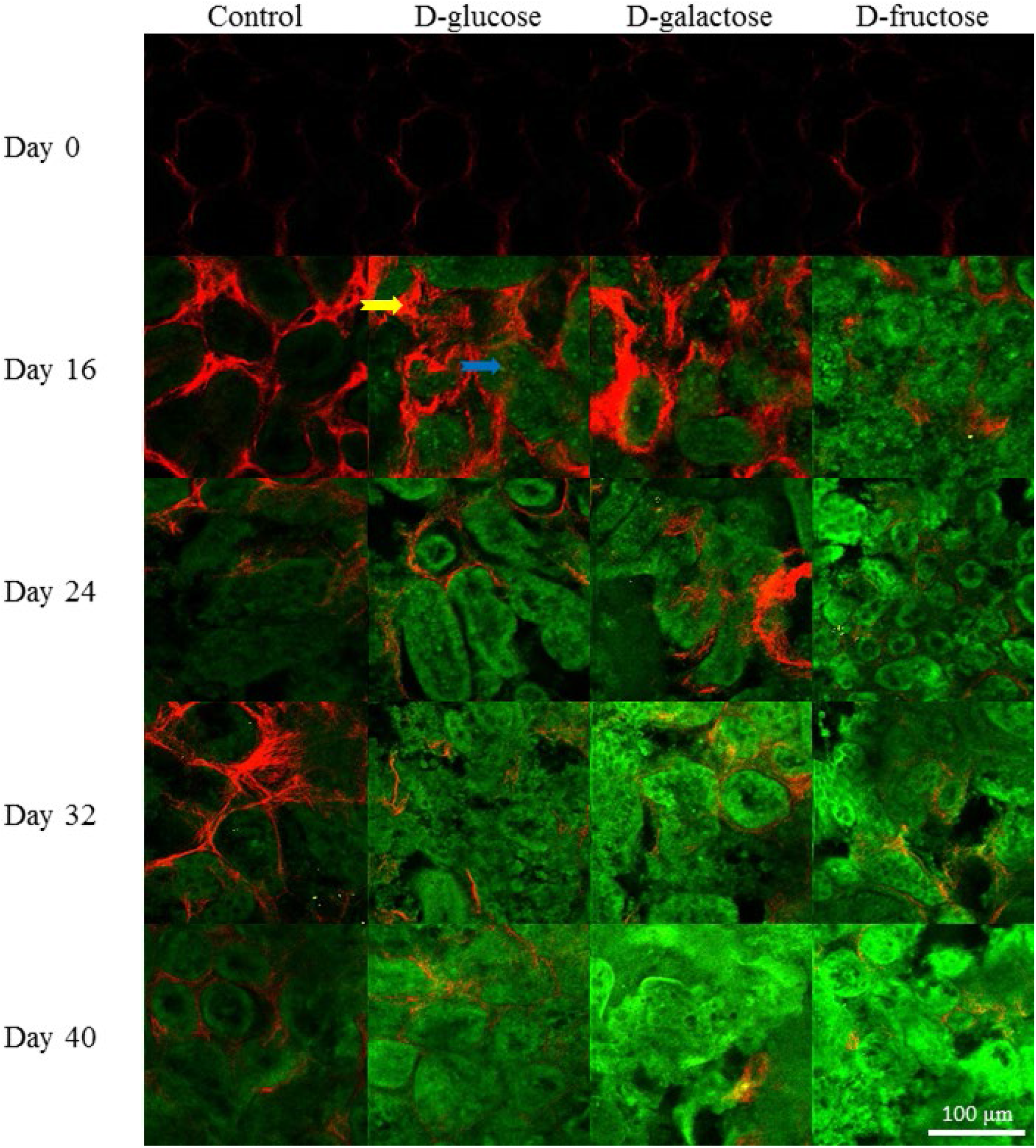
Autofluorescence and SHG imaging of porcine renal cortex in control, D-glucose, D-galactose and D-fructose solutions. Glomerular and tubular cells became strongly autofluorescent under glycation. Collagen tissue (yellow arrow) is indicated by SHG signal. Red: SHG. Green: autofluorescence. Scale bar is 100 μm.

**Figure 4.**
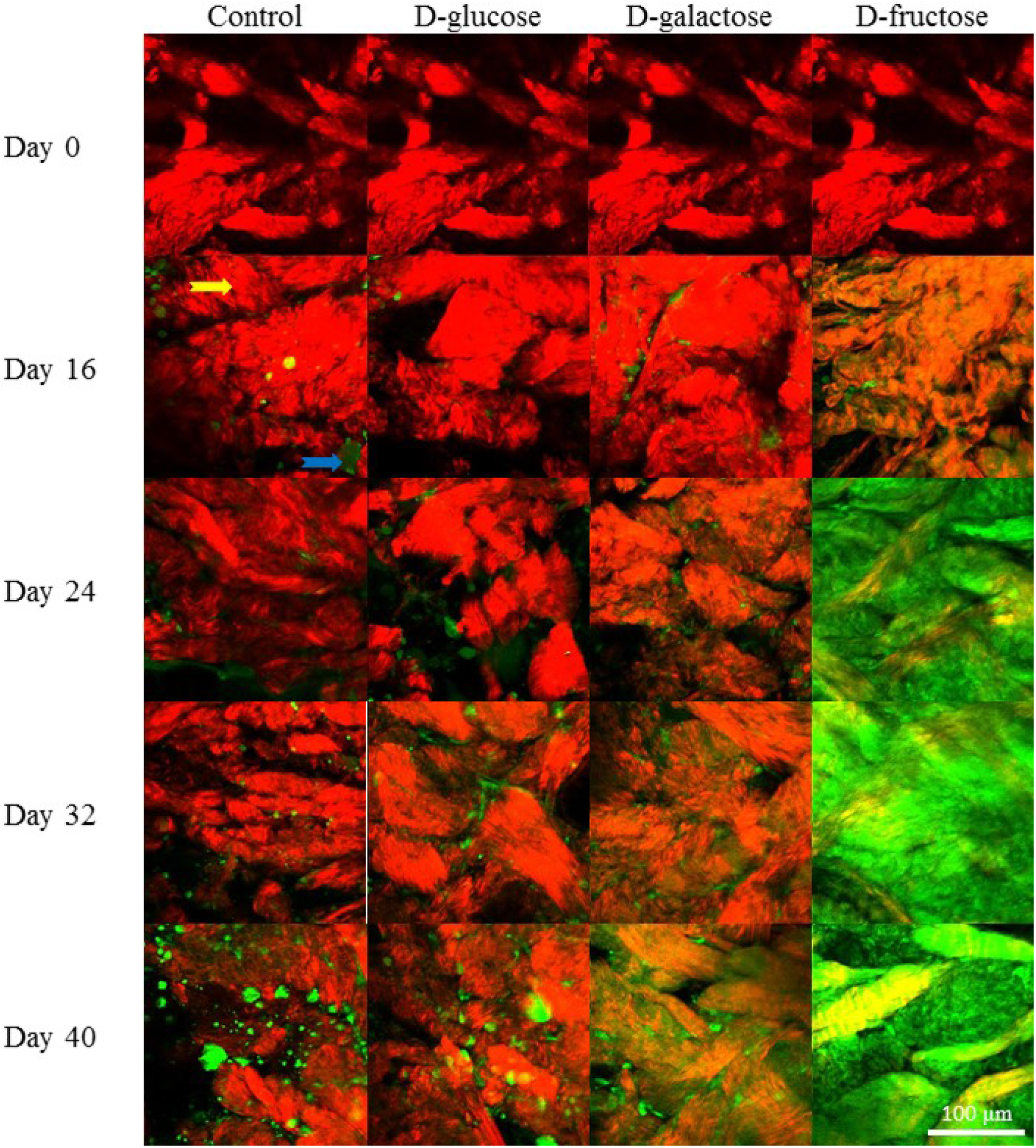
Autofluorescnece of dermic fAGE with temporal in control, D-glucose, D-galactose and D-fructose solutions. Collagen (yellow arrow), elastin, and dermal cells (blue arrow) are the main components in dermis sample. Red: SHG. Green: autofluorescence. Scale bar is 100 μm.

**Figure 5.**
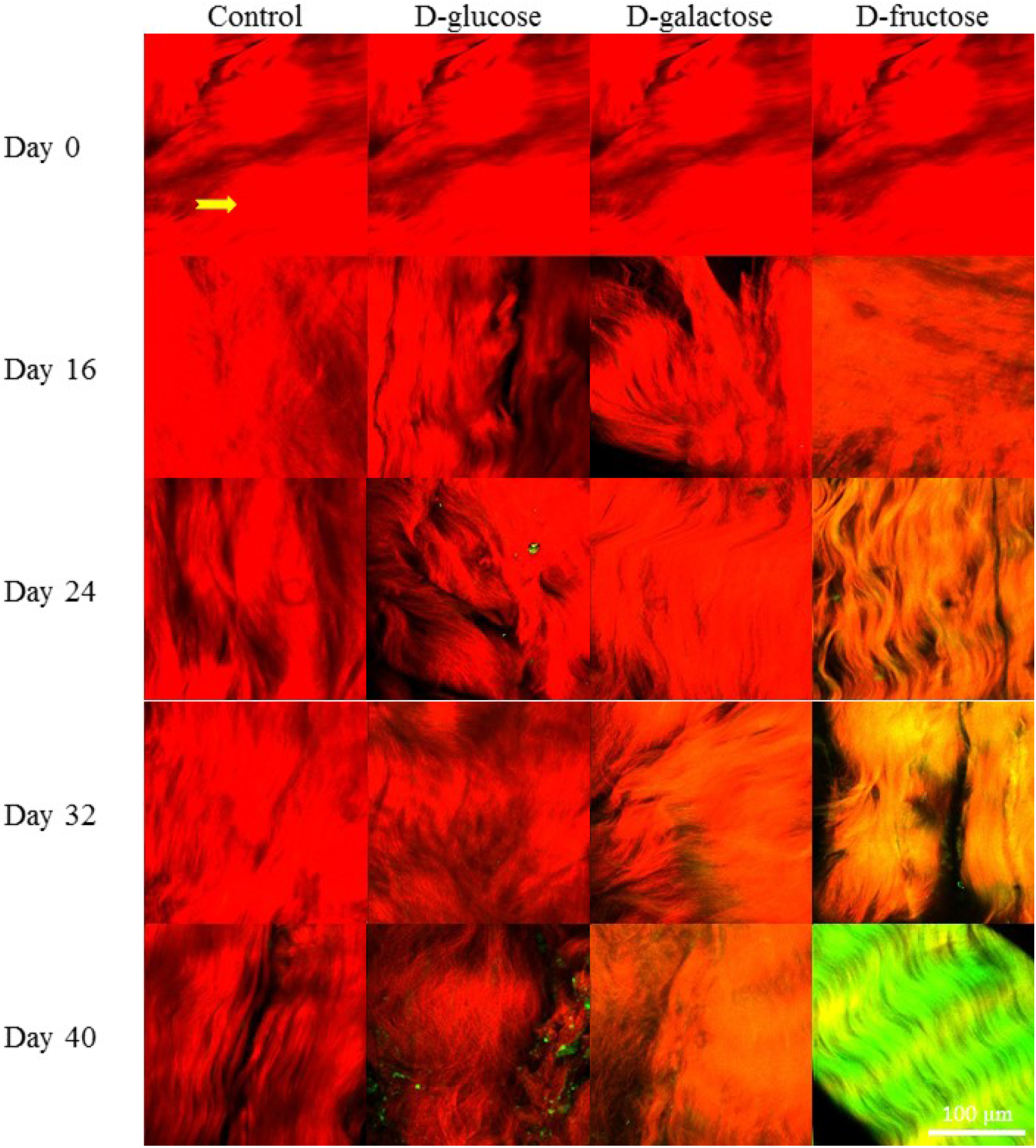
Autofluorescence of tendon with temporal in control, D-glucose, D-galactose and D-fructose solutions. The main component of tendon is collagen (yellow arrow). The increase of autofluorescence signifies the production of fAGEs. Red: SHG. Green: autofluorescence. Scale bar is 100 μm.

In Fig. 6, we analyzed the collagen autofluorescence along with the autofluorescence of other tissue components that do not generate SHG signals. Since different tissues may respond differently to monosaccharide-induced glycation, GUN Octave 5.1.0.0 was used to separate the images to determine the fAGE formation in collagen and non-collagen tissues. Specifically, collagen images were used as masks to separate the autofluorescence signals into those from collagen and those from other tissue constituents. In this manner, we determined the glycation rates of collagen in different tissues: aorta (Fig. 6(A)), cornea (Fig. 6(B)), kidney (Fig. 6(C)), dermis (Fig. 6(D)), and tendon (Fig. 6(E)). Subsequently, we determined the slope of the temporal dependence of collagen autofluorescence curves in these tissues by linearly fitting the autofluorescence intensity induced by the monosaccharide solutions as a function of time. The slopes, indicative of the rate of glycation are shown in Table 1. Our results show that D-fructose caused the fastest glycation in the selected tissue as compared to D-galactose and D-glucose. Specifically, the rates of collagen glycation in cornea, tendon, and dermis from D-glucose are between 12-14% of that due to D-fructose whereas in the aorta and kidney the same ratio is found to be 48% and 65%, respectively. On the other hand, Table 1 shows that D-galactose induced glycation rates in the cornea, tendon, aorta, dermis, and kidney at 58%, 25%, 55%, 32%, and 87%, respectively, in comparison to that due to D-fructose.

**Figure 6.**
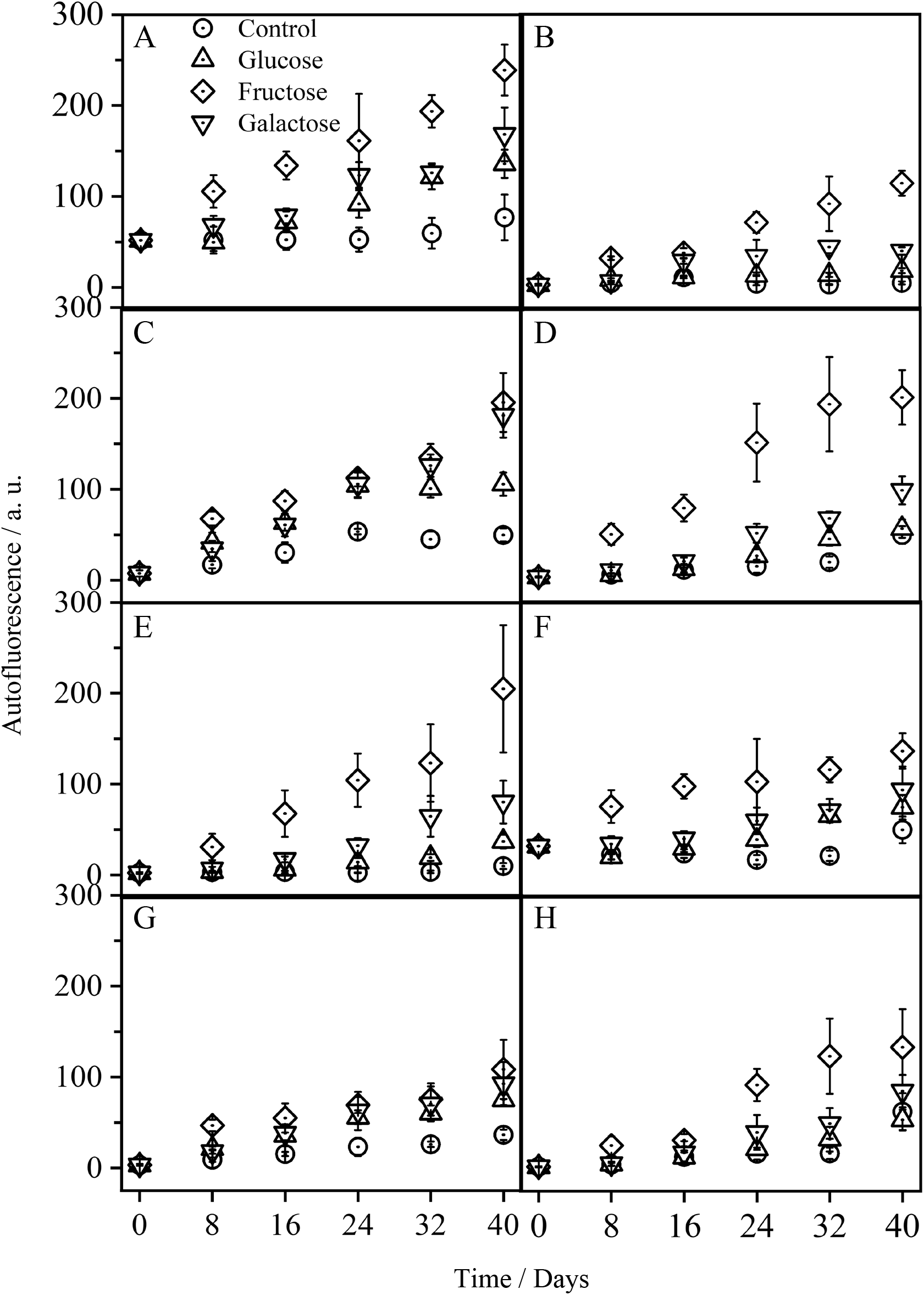
Temporal dependence of TPAF intensity in glycated collagen of (A) aorta, (B) cornea, (C) kidney, (D) dermis, (E) tendon, TPAF intensity of elastin in (F) aorta, (G) dermis and (H) glomerular/tubular cells of kidney.

**Table 1.**
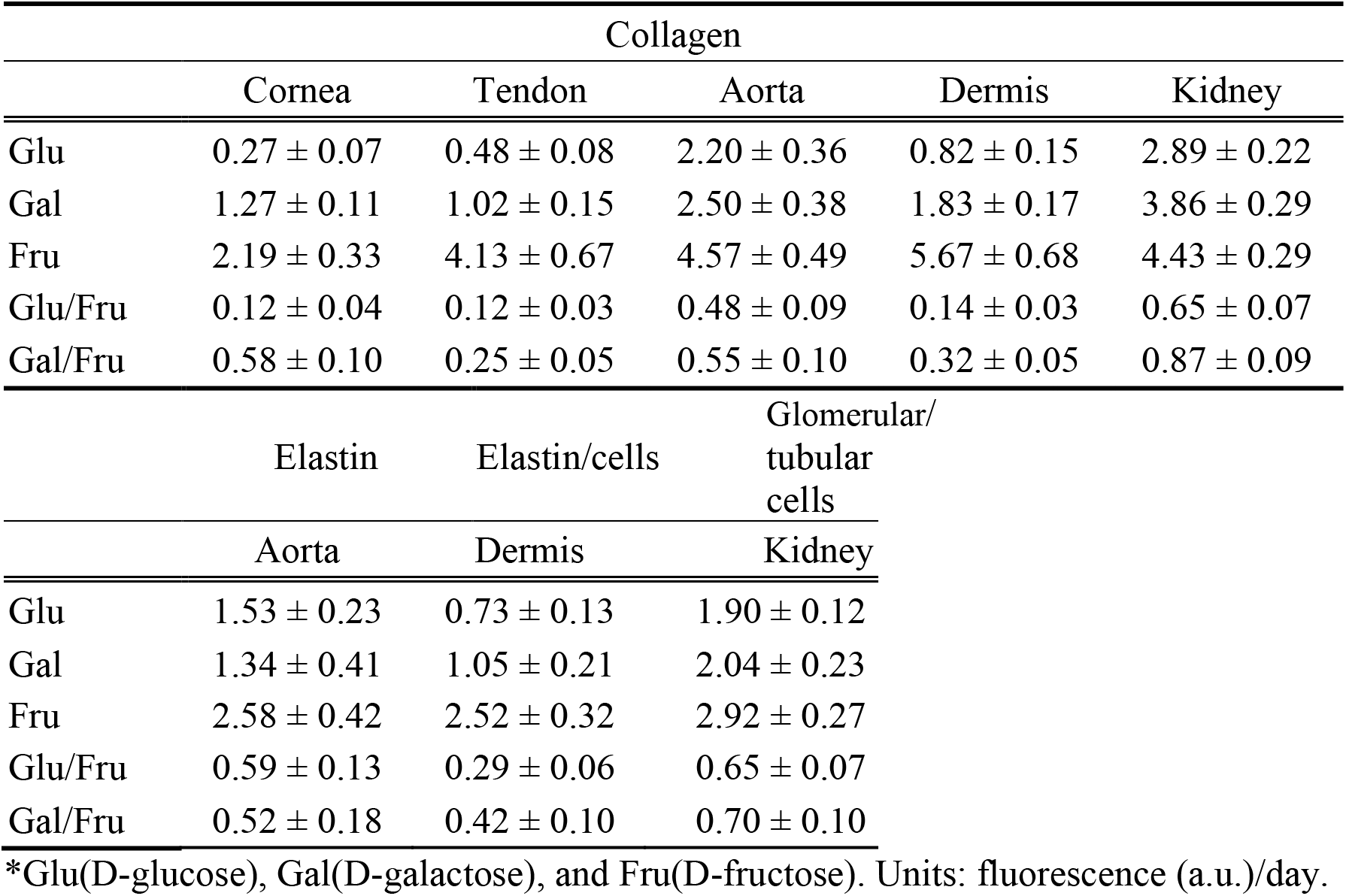
Glycation rate of collagen, elastic fiber, and glomerular/tubular cells as measured by the rate of autofluorescence increase in D-glucose, D-galactose, and D-fructose solutions.

In addition to analyzing the fAGE formation in collagen, we also analyzed the temporal dependence of autofluorescence in elastic fibers and cells. As shown, the autofluorescence intensity of elastic fibers in the aorta and dermis both increased with glycation time (Figs. 6 (F) and (G)). The same observation is also observed in the autofluorescence signal of glomerular/tubular cells in the renal cortex (Fig. 6(H)). As with collagen glycation, D-fructose resulted in the highest glycation in elastic fibers and cells. In the aorta, D-glucose and D-galactose caused the glycation rate of elastic fibers to be just 59% and 52% that of D-fructose. In dermis samples, D-glucose and D-galactose resulted in the glycation rate of elastic fibers/cells to be about 29% and 42% that of fructose. However, in glomerular/tubular cells in the renal cortex, glycation rates of D-glucose and D-galactose are 65% and 70% that of D-fructose. Also, note in the aorta and dermis, the glycation rate of collagen was found to be higher than that of the non-collagen tissue components (elastic fibers/cells). In addition to showing that D-fructose is most efficient in inducing glycation in all the tissue types we studied, Table 1 also suggests that for the same molecular type (collagen), the microscopic structural organization can affect glycation rate. Specifically, it is well-known that collagen in cornea and tendon form compact structures in three dimensions. The lower glycation rates of these two tissues suggest that a tightly organized collagen matrix hinder the formation of fAGEs. Finally, comparing the glycation rates of collagen in the renal cortex to that of glomerular/tubular cells, the higher glycation rate of collagen (4.43± 0.29 fluorescence (a.u.)/day as compared to 2.92 ± 0.27 fluorescence (a.u.)/day) suggest that collagen is a more sensitive target for detecting the formation of fAGEs in the renal cortex.

## Conclusion

In this study, we compared the rates of tissue glycation in porcine aorta, cornea, dermis, tendon, and renal cortex in D-glucose, D-galactose, and D-fructose *in vitro*. The tissues were processed and immersed in solutions containing the three monosaccharides for up to 40 days. In all tissue types, the rate of fAGE formation was the highest in D-fructose, followed by D-galactose, and then D-glucose. Compared to non-collagen tissue constituents such as elastic fibers and cells, the rate of tissue glycation was consistently higher in collagen, suggesting that collagen is a more sensitive target for fAGE formation. However, we also found that collagen in different tissues exhibits different rates of fAGE formation, with slower rates observed in tightly packed tissues such as cornea and tendon. Our study suggests that for fAGE to be developed into a long-term glycemic biomarker, loosely organized collagen that are located in the proximity of vasculature may be the best targets.

## Acknowledgment

This work was supported by the Ministry of Science and Technology, Taiwan, Republic of China (MOST 107-2112-M-002-023-MY3)

